# Functional abnormalities in the cerebello-thalamic pathways in an animal model of dystonia

**DOI:** 10.1101/2020.01.29.926170

**Authors:** Elena Laura Margarint, Hind Baba Aïssa, Andrés Pablo Varani, Romain Sala, Fabien Menardy, Assunta Pelosi, Denis Hervé, Clément Léna, Daniela Popa

## Abstract

Dystonia is often associated with functional alterations in the cerebello-thalamic pathways, which have been proposed to contribute to the disorder by propagating pathological firing patterns to the forebrain. Here, we examined the function of the cerebello-thalamic pathways in a model of DYT25 dystonia, mice carrying a heterozygous invalidation of *Gnal* gene which notably disrupts striatal function, exhibiting dystonic movements and postures following systemic or striatal administration of oxotremorine. Theta-burst optogenetic stimulations of the cerebellar nuclei evoked a potentiation of the responses to cerebellar stimulations in the thalamus and motor cortex in WT mice, without evident motor function disruption. In contrast, theta burst stimulations evoked a depression of these responses only in dystonia-manifesting *Gnal*+/− mice after oxotremorine administration, decreased the disabling dystonia attacks, and increased normal active wake behaviour in *Gnal*+/− mice. The cerebellum could thus offer a gateway for a corrective treatment of motor impairments in dystonia including striatal dysfunction.

**One sentence summary:** A mouse model of DYT25 dystonia, carrying a *Gnal* mutation disrupting striatal neurotransmission, exhibits anomalous cerebello-thalamic plasticity in the non-manifesting state, but theta-burst cerebellar stimulations during cholinergic-induced dystonia depress the cerebello-thalamic transmission and reduce the severity of the motor symptoms.

## Introduction

The purposeful motion of our body is central in the human activity, and neurological diseases altering motor function represent major clinical issues. Dystonias are characterized by involuntary muscle contractions that induce abnormal twisted positions and postures, or cause patterned or stereotyped movements (Albanese et al., 2013). Many primary dystonias have a hereditary component (Breakefield et al., 2008), but the penetrance of genetic forms of dystonia is variable and even in patients carrying identical mutation, symptoms vary in severity, age of onset, focal or generalized localization or progression (Fuchs and Ozelius, 2013; Dufke et al., 2014). This intrinsic variability in dystonia suggests the existence of interacting mechanisms that synergize or cancel out at different levels of the motor centres to determine symptom onset (Hendrix and Vitek, 2012; Prudente et al., 2014; Kaji et al., 2017).

Dysfunctions of the cerebellum and cerebello-cortical pathways have been implicated in dystonia (Neychev et al., 2011; Lehéricy et al., 2013). Patients with various forms of dystonia exhibit structural and functional alterations of the cerebellum as well as abnormalities in the cerebello-thalamic connections (Lehéricy et al., 2013). The relevance of the cerebellum in dystonia has also been demonstrated in animal models (Neychev et al., 2011). In three genetic models of dystonia *(dt* rat, tottering mouse and mouse with invalidation of type 1 inositol triphosphate receptor in the cerebellum/brainstem), the removal of the cerebellum, or only of the cerebellar Purkinje neurons or deep cerebellar nuclei (DCN), abolishes dystonic movements (LeDoux et al., 1993; LeDoux et al., 1995; Campbell et al., 1999; Neychev et al., 2008; Hensch et al., 2013). The down-regulation of the DYT1 gene, Torsin A, in the cerebellum is sufficient to trigger dystonia (Fremont et al., 2017). Neurons of the cerebellar nuclei can rapidly excite the dorsolateral striatum through a di-synaptic pathway with a relay in the centro-lateral (CL) nucleus of the thalamus (Chen et al., 2014). Aberrant cerebellar activity can cause dystonia by dynamically forcing a pathological state via the cerebello-thalamo-striatal pathway (Calderon et al., 2011). Structural defects in the cerebello-thalamo-cortical pathway have been identified in a mouse model of DYT1 dystonia (Ulug et al., 2011), but they have been proposed to provide a protective effect and limit the penetrance of the disease in patients (Argyelan et al., 2009), consistent with the view that dystonias are brain motor network disorders, where dysfunction of one node results in dysfunctions/adaptations in the others.

Dystonia also derive from basal ganglia dysfunction. One example is the adult-onset DYT25 dystonia (Fuchs et al., 2013; Vemula et al., 2013; Pelosi et al., 2017), caused by loss-of-function mutations of the *GNAL* gene encoding Gα_olf_, a G protein stimulating adenylate cyclase activity in the striatum. *Gnal* haplo-deficiency in mice mimics the genetic alterations discovered in DYT25 dystonic patients (Pelosi et al., 2017). *Gnal* haplo-deficiency reduces striatal cAMP production and disrupts striatal functions but the mice are asymptomatic in terms of dystonia. Dystonic symptoms were generated by injections of a muscarinic cholinergic agonist (oxotremorine M) systemically or in the striatum, but not in the cerebellum, indicating that an increase in striatal cholinergic tone is critical to the onset of disorder (Pelosi et al., 2017). The present study is aimed at examining the role of cerebello-thalamic tract and its plasticity in this model.

To study the functional connectivity in the cerebello-thalamic tract, we combined optogenetic stimulations in the cerebellar dentate nucleus with recordings in the thalamus and motor cortex, and with behavioural measures. Moreover, we probed the plasticity induced by cerebellar stimulations using theta-burst stimulation protocols. Indeed, transcranial theta-burst stimulations are used to treat various motor and non-motor disorders (Suppa et al., 2016; Jannati et al., 2017). When administered on the cerebellar cortex, they induce lasting changes of motor cortex excitability (Popa et al., 2010; Gallea et al., 2013). Here, we investigated the behavioural and neurophysiological impacts of dentate nucleus (DN) low frequency or theta-burst frequency optogenetic stimulation on the centro-lateral (CL), ventro-anterior-lateral (VAL) thalamus, and motor (M1) cortex in both the pre-symptomatic (saline injection), and symptomatic states (oxotremorine injection), of *Gnal*+/− animal model of dystonia.

## Results

### *Gnal*+/− mice do not have a constitutive locomotor impairment

Young 3 to 7-month-old mice *Gnal*+/− mice are asymptomatic (Pelosi et al., 2017) and to further verify that they do not exhibit constitutive motor deficit we performed a set of locomotor experiments including vertical pole, horizontal bar, grid test, fixed speed rotarod, gait test and an open-field test (Figure 1). Overall, we did not observe major motor deficits; however, mild effects were noted when the analysis was performed separately on males and females as reported below. A total of 16 *Gnal*+/− mice were studied, 12 males and 4 females. Regarding the wild-type (WT) mice, 19 mice were studied, 11 males and 8 females.

**Figure 1.**
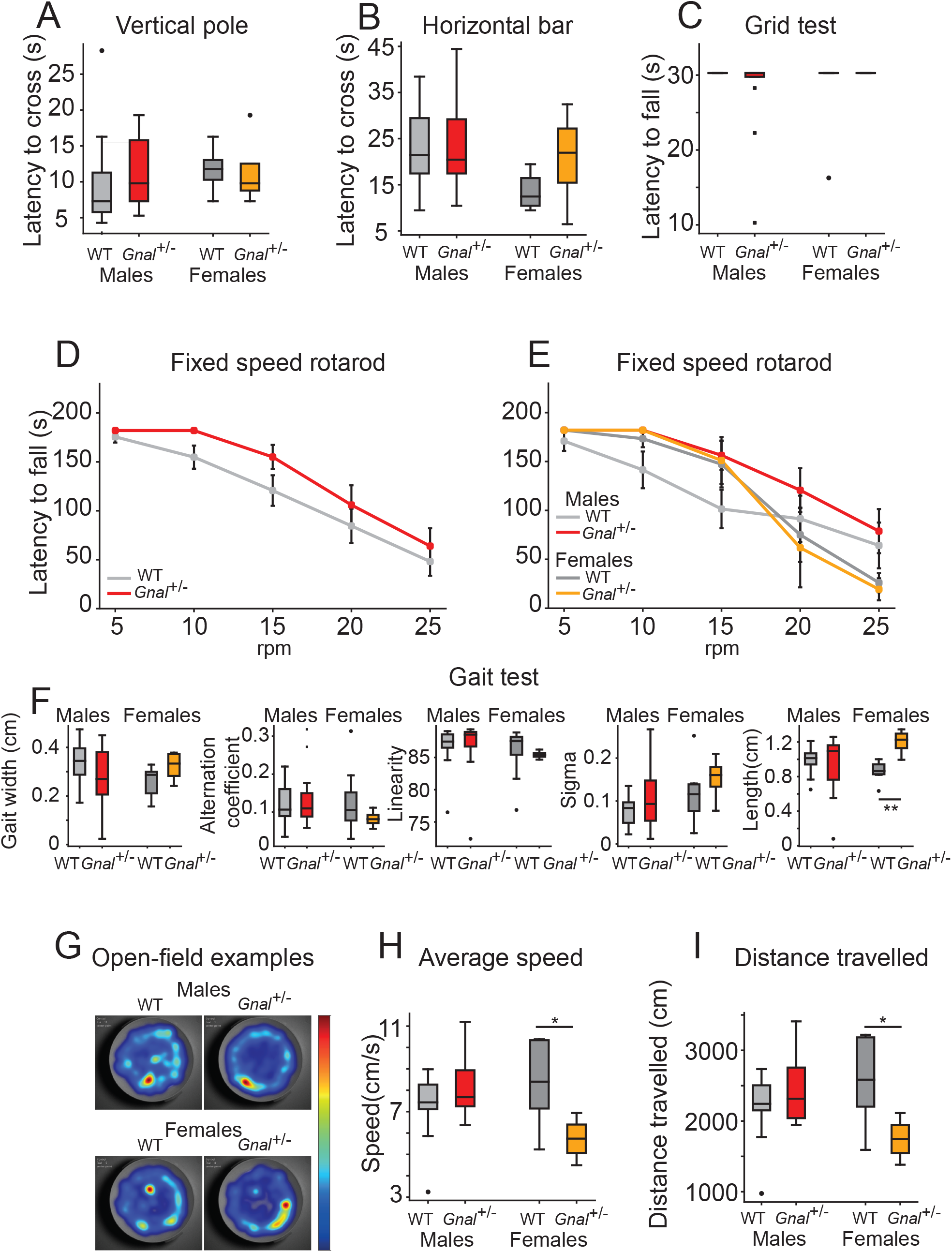
Locomotor activity and motor coordination in *Gnal*+/− mice. **(A)** Latency to cross the vertical pole; Mann-Whitney test. **(B)** Latency to cross the horizontal bar; Mann-Whitney test. **(C)** Latency to fall during the grid test with a 30s cut-off; Mann-Whitney test. **(D)** Latency to fall during the fixed speed rotarod test; Repeated measure ANOVA. **(E)** Latency to fall during the fixed speed rotarod test with groups separated by gender and genotype; Welch corrected Student’s t test for each speed step, corrected for multitests using Bonferroni’s method. **(F)** Gait width, alternation coefficient, movement linearity, sigma and stride length during the gait test; Mann-Whitney test *p<0.05; **p<0.01 **(G)** Example of heatmaps showing the position of mice during open-field sessions. Average speed **(H)** and total distance travelled **(I)** during an open-field session; Mann-Whitney test *p<0.05; **p<0.01.

We observed that the motor performance of *Gnal*+/− mice was not impaired in the vertical pole test (Mann-Whitney test; p = 0.176 for males; p = 0.246 for females), horizontal bar test (Mann-Whitney test; p = 0.487 for males; p = 0.134 for females) and grid test (Mann-Whitney test; p = 0.164 for males; p = 0.298 for females).

We also performed a fixed speed rotarod test to test the motor coordination. We did not find significant differences between *Gnal*+/− and WT mice (Repeated measure ANOVA F(1,4)=7.348 p = 0.053), nor when comparing males and females for each speed steps (Student’s T test Welsch corrected, with a Bonferroni correction for multitests, p= 0.171 for both males and females).

We performed the gait test and different parameters were computed. We were not able to find significant differences in gait width (Mann-Whitney test; p = 0.154 for males; p = 0.074 for females), alternation coefficient (Mann-Whitney test; p = 0.391 for males; p = 0.175 for females), linear movement (Mann-Whitney test; p = 0.221 for males; p = 0.101 for females) or sigma (Mann-Whitney test; p = 0.488 for males; p = 0.134 for females). However, the length of stride was significantly longer in *Gnal*+/− females compared to female WT mice (Mann-Whitney test; p = 0.007 for females; p = 0.322 for males).

We also studied both the average locomotor speed and the total distance travelled during an open-field session. We observed no significant differences when comparing *Gnal*+/− and WT males (Mann-Whitney test; p = 0.141 for the average speed; p = 0.203 for the distance travelled). However, we observed that both average speed and distance travelled were lower in females *Gnal*+/− compared to females WT (Mann-Whitney test; p = 0.037 for the average speed; p = 0.037 for the distance). In conclusion, motor activity and motor coordination are not dramatically impaired in young 3 to 7-month-old mice *Gnal*+/− mice as previously described (Pelosi et al., 2017); however, mild differences were observed in females only, raising the possibility of higher susceptibility in female mice.

### Thalamic and cortical activity, and coupling of the cerebellum to forebrain motor circuits in *Gnal*+/− mice

The thalamus is the main gateway to the motor cortex and striatum, which are, together with the cerebellum, the main structures involved in dystonia. To examine the impact of the reduction of *Gnal* expression in the thalamus, we first examined the changes in neuronal activity in the motor thalamus (VAL), the striatum-projecting thalamus (CL) and the motor cortex (M1) in freely moving mice (Figure 2, Table 1). Saline injections in *Gnal*+/− mice did not induce abnormal motor patterns, consistent with previous observations (Pelosi et al., 2017). In these mice, we found no difference as compared to saline-injected wild type (WT) mice in the baseline firing rate of thalamic or cortical neurons (Figure 2E, ANOVA repeated measure VAL: F(1,16)=0.19, p=0.67, CL: F(1,21)=0.21, p=0.65, M1: F(1,23)=0.033, p=0.86; no difference was found when males and females were separated).

**Table 1.**
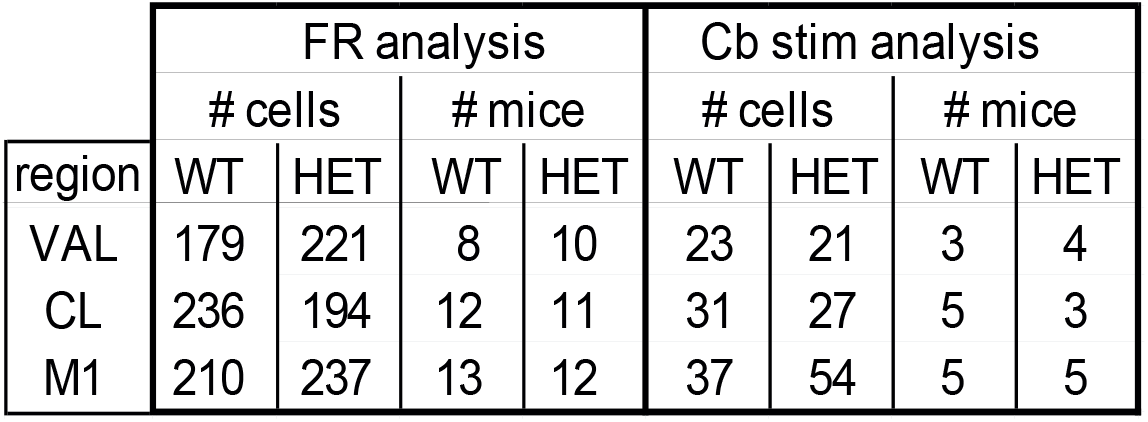
Number of cells used in the analysis of baseline firing rate (Figure 2E) and of responses to cerebellar stimulation (Figure 3C,D,G,H).

| region | FR analysis |  |  |  | Cb stim analysis |  |  |  |
| --- | --- | --- | --- | --- | --- | --- | --- | --- |
|  | # cells |  | # mice |  | # cells |  | # mice |  |
|  | WT | HET | WT | HET | WT | HET | WT | HET |
| VAL | 179 | 221 | 8 | 10 | 23 | 21 | 3 | 4 |
| CL | 236 | 194 | 12 | 11 | 31 | 27 | 5 | 3 |
| M1 | 210 | 237 | 13 | 12 | 37 | 54 | 5 | 5 |

**Figure 2.**
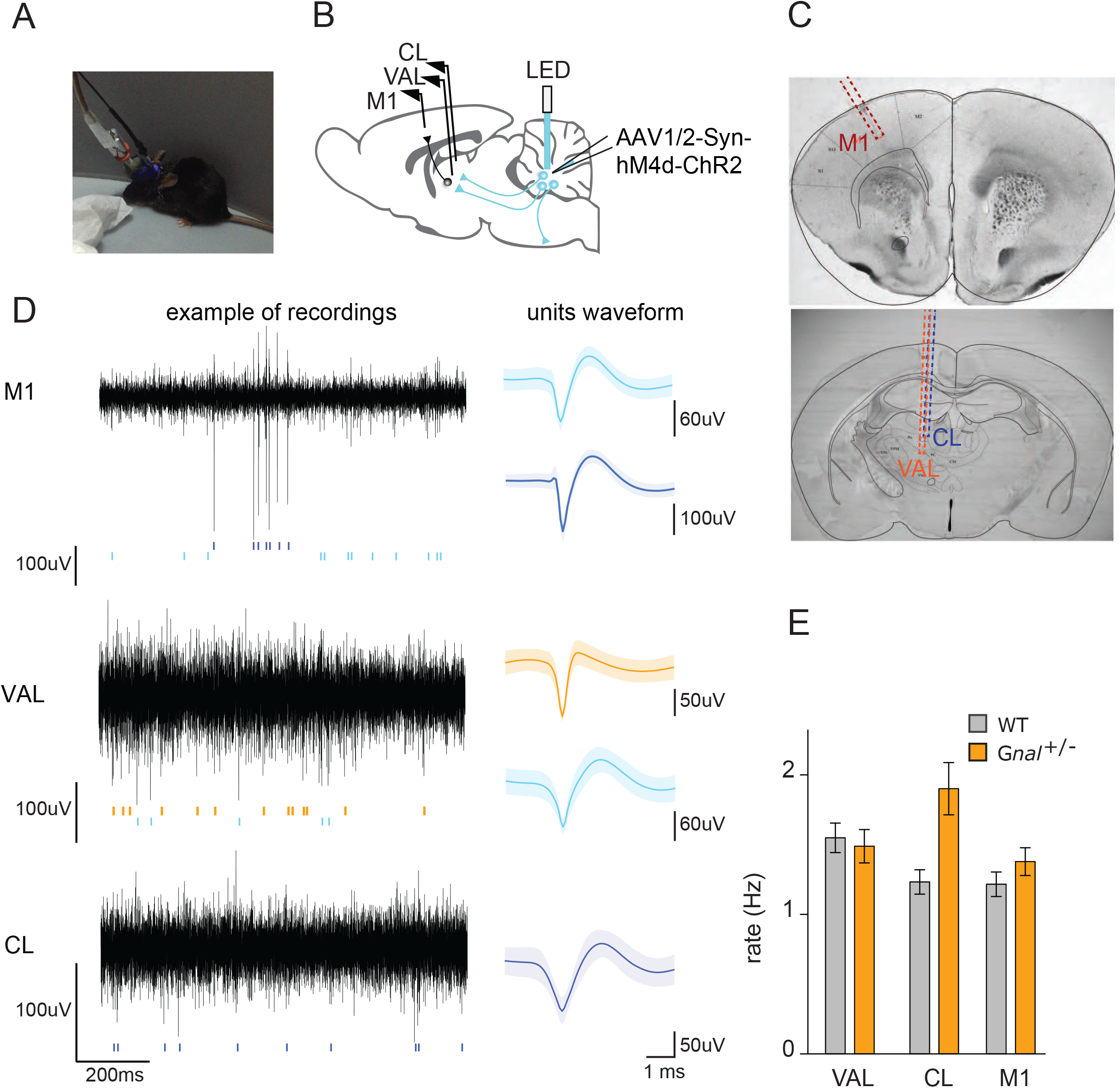
Electrophysiology recordings of ventro-lateral VAL and centro-lateral CL thalamic nuclei and (M1) motor cortex. **(A)** Picture of a freely moving mouse during a recording coupled with optogenetic DN stimulation. **(B)** Schematic of experimental design. Guide cannulas with bundles of electrodes were inserted in left M1 (motor cortex), CL (centro-lateral thalamus), VAL (ventro-lateral thalamus); an optic fibre was inserted in the right DN 3 weeks after AAV virus was injected. **(C)** Histology section showing the implantation sites. An atlas map is overlaid on the coronal sections to identify the targeted regions. **(D)** Left: excerpts of traces and spikes (colored bars); right: average ± SEM waveforms of VAL, CL and M1 neurons. **(E)** Average firing rate in freely moving WT and *Gnal*+/− mice.

To probe the functional drive of the cerebellum on these regions, we expressed channelrhodopsin 2 (ChR2) into DN neurons using an AAV viral strategy (AAV2/1.hSyn.ChR2(H134R)-eYFP.WPRE.hGH), and stimulated these neurons through an optical fibre implanted above the DN (Figure 2B, 2). We first applied simple low-frequency deep cerebellar stimulations (100ms, 0.25Hz, 0.9mW at the fibre entry), and we found an overall increase in firing rate of about 5 Hz during the stimulation in all recorded regions in WT and in *Gnal*+/− saline-injected mice. We did not detect any significant difference between the genotypes (Figure 3C, ANOVA with repeated measure in mice: CL: F(1,6)=1.057, p=0.343), VAL: F(1,5)=1.466, p=0.28, M1: F(1,8)=0.0395, p=0.847). We then performed oxotremorine M administration (0.1 mg/kg, i.p.) and observed an increase in the evoked responses to cerebellar stimulation in all the structures studied in both the WT and *Gnal*+/− genotypes (Figure 3G, Table 2).

**Table 2.**
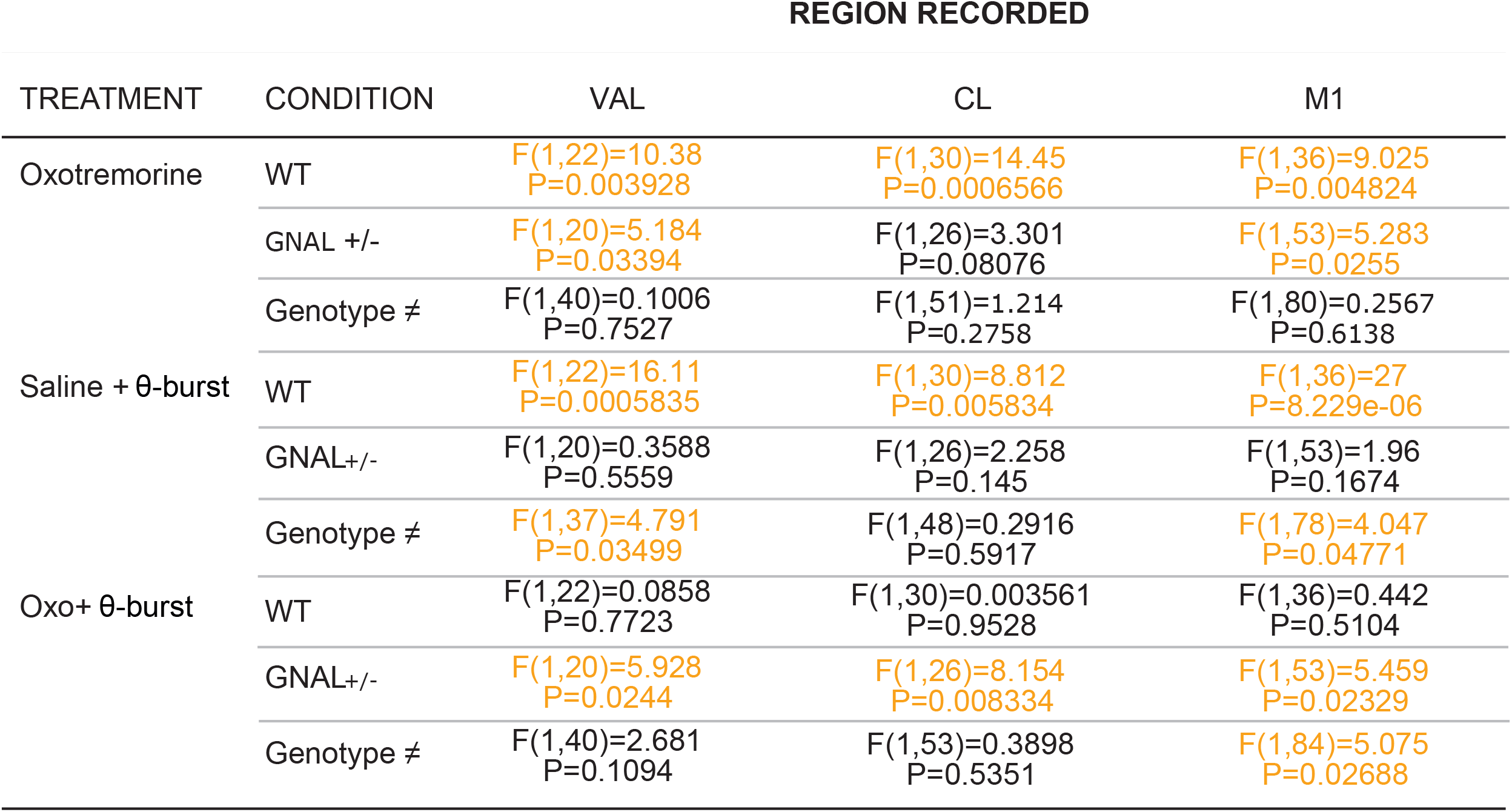
Repeated measure ANOVA for the increase in firing rate induced by low-frequency DN stimulations in the VAL and CL thalamus, and motor cortex M1 in saline/oxotremorine conditions and before/after theta-burst stimulations.

**Figure 3.**
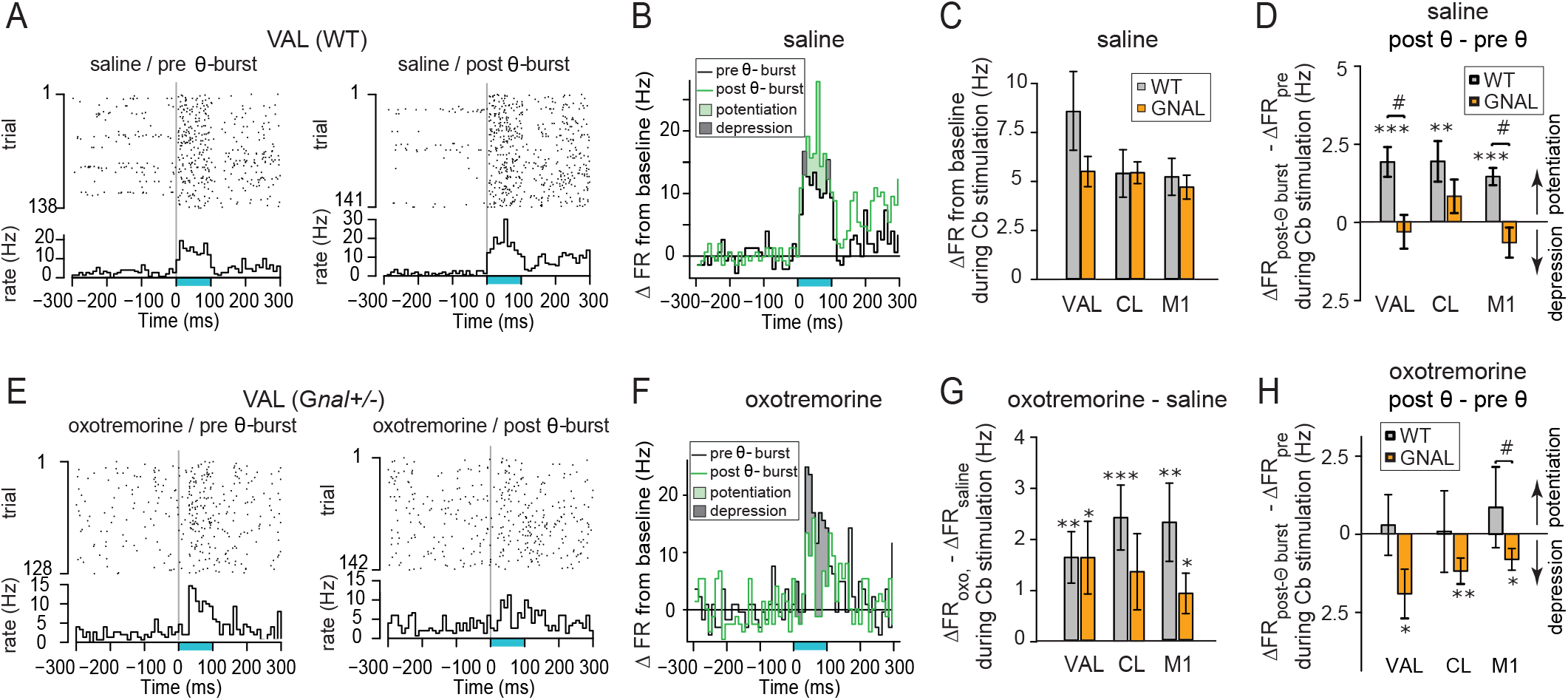
Effect of low-frequency and theta-burst cerebellar stimulations on the firing rate in saline and oxotremorine conditions in VAL, CL and M1 of WT and *Gnal+/−* mice. On the first day, we examined the effects of low frequency opto-stimulations (blue line, 100ms, 0.25Hz) on the firing rate of neurons in the various brain regions after saline and oxotremorine administrations. On the following day, the effects of the same low frequency stimulation were compared before and after θ-burst stimulations (20ms, 8,33Hz, applied for 2 × 40s with a 2 min pause in between) after saline and oxotremorine administrations. **(A)** Raster and peristimulus time histogram (PSTH, bin 10 ms) for one VAL neuron from a WT mouse before (left) and after (right) theta-burst stimulation in saline condition; the PSTH are normalized to express a firing rate (see Methods). **(B)** Overlay of the PSTHs from the panel A; the difference between the histograms is filled to evidence the potentiation or depression of the responses. In this case potentiation prevails. See Table 1 for number of cells/mice used. **(C)** Increase in firing rate (relative to the baseline firing rate) induced by 100 ms DN nucleus stimulations in WT and *Gnal*+/− animals. **(D)** Impact of theta-burst stimulations administered in saline condition on the response to 100 ms DN stimulation. Note the absence of potentiation in *Gnal+/−* mice. **(E) (F)** same as (A) and (B) respectively, for a VAL neuron from a *Gnal*+/− mouse before and after theta-burst stimulation in the oxotremorine condition. **(G)** effect of oxotremorine administration on the response to cerebellar stimulations. **(H)** Impact of theta-burst stimulations on the response to 100 ms DN stimulation (same as (D)) in the oxotremorine condition. See Table 2 for corresponding repeated measures ANOVAs. *p<0.05, **p<0.01, ***p<0.001 paired difference between conditions for the cells, ^#^p<0.05 difference between *Gnal*+/− and WT mice. Error bars represent SEM.

To examine the plasticity of the cerebello-thalamic pathways, we then studied the impact of optogenetic theta-burst stimulations (given the limited number of cells, males and females were not studied separately). In the saline condition, the response to low-frequency stimulations was significantly increased in CL, VAL (Figure 3A,B) and M1 after theta-burst stimulations in the wild type mice, while no significant change was found in the *Gnal*+/− mice (Figure 3D, Table 1 & 2). The difference between wild type and Gnal+/− mice was clearly significant in VAL and M1 (Figure 3D, Table 1 & 2). In contrast, following oxotremorine administration, the responses to cerebellar low-frequency stimulations were not changed in wild type mice after theta-burst stimulations, while they were decreased in *Gnal*+/− mice in all the structures considered: CL, VAL (Figure 3E,F) and M1 (Figure 3H, Table 1 & 2), suggesting a weaker entrainment of the motor circuit by the cerebellum during dystonic-like attacks induced by oxotremorine in *Gnal*+/− mice. The increase (potentiation) (Figure 3A,B) or decrease (depression) (Figure 3E,F) in firing rate were observed throughout the duration of the stimulation (Figure 3B,F).

### Beneficial effect on motor activity of the optogenetic stimulation in the cerebellar dentate nucleus of *Gnal*+/− mice

Taking advantage of chronic implanted animals, we then evaluated the dystonic behaviour in *Gnal*+/− compared to WT mice using an abnormal motor score (Fremont et al., 2017; Pelosi et al., 2017; White and Sillitoe, 2017). Saline-injected *Gnal*+/− and WT mice showed no signs of dystonia; in contrast, the muscarinic cholinergic agonist, oxotremorine M (0.1 mg/kg), consistently induced abnormal postures such as extension of hind limbs from the body axis for >10 s, sustained hunched posture with little movements, slow walking with increased hind limb gait yielding maximal dystonia scores in all implanted *Gnal*+/− mice and only mild motor signs in wild type mice (Figure 4A,B). These observations are consistent with the previous observations in *Gnal*+/− mice (Pelosi et al., 2017) and indicate that the chronic implantation did not impact the development of dystonic-type motor abnormalities.

**Figure 4.**
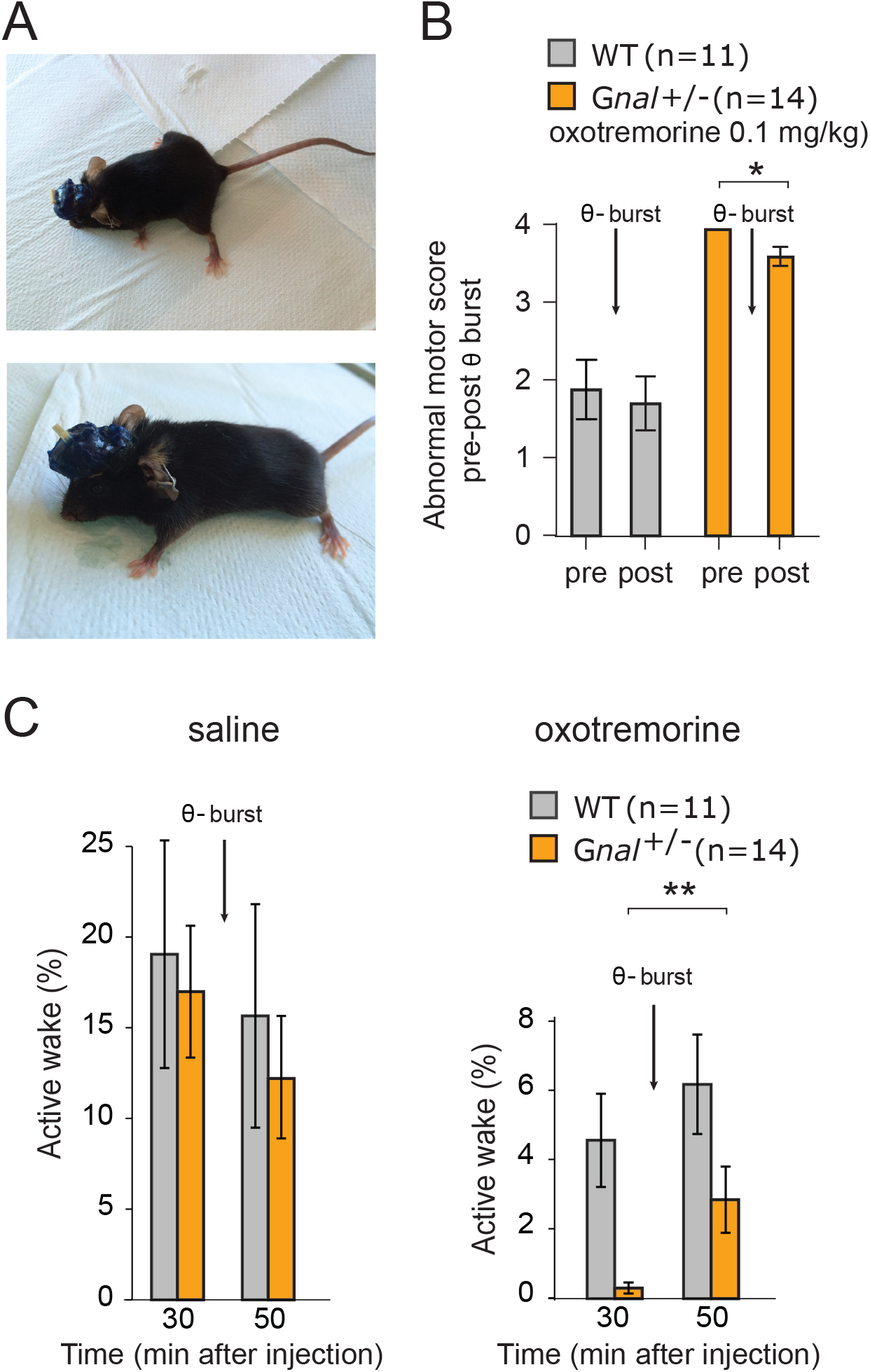
Motor behavior of *Gnal*+/− (orange) and WT mice (grey). **(A)** Examples of dystonic postures in *Gnal*+/− mice following oxotremorine M administration. **(B)** Average dystonia scores in *Gnal*+/− and WT mice following oxotremorine M administration before and after dentate nucleus (DN) theta burst stimulation **(C)** Change of average active wakefulness percentage after one session of DN theta-burst stimulation in *Gnal*+/− and WT mice. Data were analysed by repeated measure Student t test. **p<0.01 difference between pre and post theta-burst stimulations. Error bars represent SEM.

To test the impact of cerebellar stimulations on abnormal movements, the evolution of dystonic-like movements and postures was evaluated before and after the DN optogenetic theta stimulations (Figure 4A,B). DN optogenetic theta-burst stimulations decreased the abnormal motor behaviour in *Gnal*+/− mice following oxotremorine M challenge (Wilcoxon matched-pairs signed rank test; P=0.0313). To confirm these observations, we also evaluated the activity of the mice by measuring the percentage of time spent moving in an open-field (“active wakefulness”) (Georgescu et al., 2018). In the saline-treated wild type mice, DN optogenetic theta burst stimulations had no significant effect (30 to 50 min after saline injection) on the active wake time. Similarly, in the saline-treated *Gnal*+/− mice, theta-burst stimulation did not significantly change the active wake time (Figure 4C). In contrast, in the oxotremorine condition, the average active wake time decreased in both wild type and *Gnal*+/− mice compared to saline, the effect being more marked in the mutant mice (Figure 4C). One session of DN optogenetic theta-burst stimulation was sufficient to significantly increase the wake time in *Gnal*+/− mice (p=0.0099) while the increase remained non-significant in wild type mice (Figure 4C). Overall, these results indicate that DN optogenetic theta-burst stimulations decrease the “dystonic” phenotype in oxotremorine-treated *Gnal*+/− mice regarding both the abnormal motor scores and locomotor activity deficit.

Overall, these results indicate a functional impairment of cerebello-thalamo-cortical pathway in *Gnal*+/− mice, which is revealed by a loss of potentiation following theta-burst stimulations in saline conditions. These stimulations decrease cerebello-thalamic connectivity during dystonic-like attacks in *Gnal*+/− mice and reduced the dystonic signs.

## Discussion

In the present study, we investigated the behaviour and functional coupling of the dentate (DN) nucleus of cerebellum to the thalamus and motor cortex M1 in basal condition and during dystonic-like state triggered pharmacologically in the *Gnal*+/− animal model. We confirmed the presence of dystonic-like movements and postures in these mutant mice after oxotremorine, a muscarinic cholinergic agonist. We found little if any change in locomotor activity or motor coordination and in baseline firing rate in the thalamus and M1 cortex between *Gnal*+/− and control mice. Theta-burst cerebellar DN stimulations induced potentiation of cerebello-thalamic pathways in wild type mice, but not in *Gnal*+/− mice suggesting a functional alteration of the cerebello-thalamic pathways in these mice. In contrast, theta-burst stimulation of cerebello-thalamic pathway induced a depression of the cerebello-thalamic connections after oxotremorine administration in G*nal*+/− mice (but not in WT mice). Such changes might prevent the propagation of the abnormal activity of the cerebellum to motor circuits in the *Gnal*+/− mice in oxotremorine condition. Indeed, theta-burst stimulation of the DN cerebellar nucleus decreased oxotremorine-induced dystonic-like motor abnormalities and increased normal active behaviour (active wakefulness) in *Gnal*+/− mice.

### Dystonia and cerebello-thalamic pathways

Despite the strength of the association between dystonia and basal ganglia (BG) demonstrated by numerous studies in both patients and animals (Köhling et al., 2004; Hallett, 2006; Fujita and Eidelberg, 2017; Simonyan et al., 2017; Huebl et al., 2018), the advances in neuroimaging approaches and electrophysiology have provided evidence that regions outside the BG are also involved. The dystonia should then be viewed as a “circuit disorder” (Lehéricy et al., 2013) spreading over a network of brain motor regions. It is now established that the cerebellum is part of this network, since compelling evidence shows that disruption of cerebellar output causes dystonia (Pizoli et al., 2002; Raike et al., 2013). Abnormal cerebellar nuclei bursting activities were observed in several animal models of dystonia such as DYT1 (Fremont et al., 2017) Tewari et al., 2017), genetic disruption of the olivo-cerebellar circuit (White and Sillitoe, 2017) or Na+/K+-ATPase α3+/D801Y mutant mice (Fremont et al., 2015; Isaksen et al., 2017). These activities are thought to rely on a pathway joining the cerebellum to the CL thalamus and striatum to propagate pathological activities (Calderon et al., 2011). This cerebello-thalamo-striatal pathway seems to be a key player in such forms of dystonia: imaging studies in DYT1 and DYT6 gene carriers and patients suggested a decreased connectivity between the cerebellum and thalamus and a disruption of cerebellum output that could play a significant role in the appearance of the dystonia (Carbon et al., 2008; Argyelan et al., 2009). If cerebellar dysfunction may disrupt basal ganglia function, the reverse has received little attention so far.

Here we studied the *Gnal*+/− mouse model of the adult-onset dystonia DYT25 (Pelosi et al., 2017). The *Gnal* gene encodes the α subunit of the heterotrimeric G-protein Golf, which is an essential relay in dopamine and adenosine transmission onto the adenylyl cyclase in the striatum. Gnal+/− mice are also characterized by abnormal responses of striatal cholinergic interneurons (Eskow Jaunarajs et al., 2019). Despite the reduced production of striatal cAMP in these mice, we found no innate motor impairment in *Gnal*+/− mice for males and only a mild difference for females, in various motor tests (vertical pole, horizontal bar, grid test, fixed speed rotarod, gait test and open-field locomotor experiments). Therefore, these mice exhibit no overt motor deficit in baseline condition, which may correspond to a ‘presymptomatic’ state. Oxotremorine injections in these mice -performed either systemically or locally into the striatum-induce strong dystonic-like syndrome compared to wild type mice(Pelosi et al., 2017). Under oxotremorine, these mice are likely representing a model of dystonia caused by striatal primary dysfunction, offering an opportunity to examine the function of the cerebello-thalamic connection in such model.

We did not find any differences in the spontaneous firing activity between wild type and *Gnal*+/− mice in CL and VAL thalamus or M1 cortex, indicating an absence of overt baseline deficits. Oxotremorine injections mildly increased the average firing rate in all structures, but there was no strong difference between wild type and *Gnal*+/− mice. Examining the activity evoked in these brain regions by cerebellar stimulation failed also to reveal overt differences, although changes in cerebello-thalamic efficiency might be concealed by the variability of parameters of the experiments (e.g. variability in ChR2 expression, in optical fiber positioning). Indeed, our plasticity experiments using theta-burst stimulations in the DN nucleus of the cerebellum revealed clear differences in the cerebello-thalamic pathway of *Gnal*+/− mice, with a lack of potentiation in the saline condition, and a depression – not observed in wild type mice-in the oxotremorine condition. These results suggest a saturation of potentiation of the cerebello-thalamic pathway of *Gnal*+/− mice, which could be reversed by stimulations in the oxotremorine condition. Such changes in plasticity and saturation are also found in the striatum in motor disorders (Calabresi et al., 2016). These anomalous plasticities might result from compensatory alterations to the striatal dysfunction induced by oxotremorine in *Gnal*+/− mice (Pelosi et al., 2017). Our results are consistent with those of White and Sillitoe (2017), who demonstrated in a model of dystonia caused by silencing the olivo-cerebellar transmission, that high frequency electrical stimulations of the interposed cerebellar nuclei decreased the dystonia-like behaviour of mice, an effect reproduced by cerebellar lidocaine inhibition consistent with an inhibitory effect of stimulations (White and Sillitoe, 2017).

### Cerebellar theta-burst stimulations and dystonia

The rationale of using theta-burst stimulation derives from human studies, where lasting changes in excitability have been shown to be induced by such stimulations (Huang et al., 2005). Transcranial cerebellar stimulations are known to affect within few milliseconds both the inhibition and excitation in the motor cortex networks (Daskalakis et al., 2004), presumably via the di-synaptic excitatory connection linking the cerebellum to the motor cortex via the ventral thalamus (Allen and Tsukahara, 1974; Iwata and Ugawa, 2005). Theta-burst protocols applied to the cerebellum have then been found to induce long-lasting changes in motor response to motor cortex stimulations (Koch et al., 2008).

Cerebellar stimulations in dystonic patients have been mostly performed in the context of focal dystonia (cervical or hand) with good results for cervical dystonia. During a sham-controlled trial on cervical dystonic patients, Koch et al. have applied two-weeks of continuous theta-burst stimulation (TBS) and transcranial magnetic stimulation (TMS) and reported a clinical improvement of 15% at the end of the trial (Koch et al., 2014). No effect was observed in hand focal dystonia (Hubsch et al., 2013; Linssen et al., 2015; França et al., 2018), but an insufficient number of repetition may be responsible for the lack of effect (Meunier et al., 2015). The stimulations are administered on the cerebellar cortex, so the site(s) of plasticity responsible for the improvement of the patient’s condition is unknown. In healthy individuals, transcranial cerebellar stimulations modulate the cortical plasticity in a paired-associative stimulation (PAS) paradigm (Hamada et al., 2012), and theta burst cerebellar stimulations may modulate the PAS remotely in time (Popa et al., 2013), indicating that the cerebellum controls cortical plasticity. Interestingly in patients with cervical dystonia, the impact of cerebellar manipulations on cortical plasticity was reversed compared to healthy controls (Popa et al., 2018). Our experiments suggest that theta-burst stimulations may also impact the cerebello-thalamic pathway, but as in cervical dystonia (Popa et al., 2018), stimulations reversing the cerebello-thalamic plasticity anomaly improve the mouse motor function, suggestive that manipulations aimed at recruiting cerebello-thalamic plasticity could be a relevant therapeutic strategy.

### Control of striatal cholinergic interneurons by thalamo-striatal inputs (BG-CB interaction)

The striatum has been shown to receive di-synaptic inputs from the cerebellum (Bostan et al., 2013), and these inputs are presumably relayed to a large part through the intralaminar thalamus CL (Ichinohe et al., 2000; Chen et al., 2014). These nuclei also send strong projections to the dorsal striatum while only modestly contributing to the innervation of the cerebral cortex (Smith et al., 2014). The principal synaptic targets of the thalamic nuclei in the striatum are the dendritic spines of the medium spiny neurons (MSNs), but they also engage cholinergic interneurons in a feed-forward regulation that differentially control the populations of MSNs expressing the D1 dopamine receptors and exerting pro-kinetic effect, and the MSNs expressing the D2 receptors and exerting an anti-kinetic effect (Ding et al., 2010). Brief stimulations of thalamic input generate in cholinergic interneurons, a burst-pause firing pattern that transiently suppresses cortical drive of the two types of MSNs and then creates a second-long period in which activity in D2-expressing MSNs is increased, producing a potent suppression of action. Since salient sensory stimuli activate intralaminar thalamic neurons, Surmerier and coll. proposed that these processes could be a key component of the physiological responses elicited in the striatum by salient sensory stimuli (Ding et al., 2010; Smith et al., 2014). These responses would produce a cessation of ongoing motor activity and later enable switching to different actions. Interestingly, in mice with the DYT1 dystonia mutation, brief stimulations of thalamo-striatal inputs, mimicking an effect of salient events, evoke abnormal responses in cholinergic interneurons, leading to an altered cortico-striatal synaptic activity of MSNs (Sciamanna et al., 2012). However, these abnormalities are probably related to a predisposition to dystonia rather than to dystonia manifestations since the animal model is essentially asymptomatic. In the *Gnal*+/− dystonia model, the dystonic state is induced by a cholinergic pharmacological activation (Pelosi et al., 2017), which likely produces a severe imbalance in the activities of the two populations of MSNs. The “anti-dystonic” effects we induced by stimulations of cerebellar output in the dentate nucleus of *Gnal*+/− mice might rely on a reduction of these alterations. However, understanding the mechanisms at play will probably require solving the complex rules of synaptic plasticity in the striatum (reviewed in Perrin and Venance, 2019).

In conclusion, our study investigates an original model of dystonia that mimics the genetic alterations discovered in DYT25 dystonic patients, a subtype of dystonia that has not yet been studied in detail. Despite the fact that the striatum is likely the primary origin of functional alterations (Pelosi et al., 2017), our study revealed abnormalities in cerebello-thalamic pathways in *Gnal*+/− mice with different plasticity properties in asymptomatic mice (in which dysfunctions are compensated) and during dystonia-like state (in response to oxotremorine). The identification of patterns of cerebellar stimulation maximizing the depression of the cerebello-thalamic pathway could thus potentially improve the strategies for therapeutic interventions in patients.

## Acknowledgments

This work was supported by Agence Nationale de Recherche to D.P. and D.H. (ANR-16-CE37-0003 Amedyst) and to C.L. (ANR-17-CE37-0009 Mopla) and by the Labex Memolife and the Institut National de la Santé et de la Recherche Médicale (France). The authors declare no competing financial interests.

## Materials & Methods

### Animals

All experiments were performed in accordance with the guidelines of the European Community Council Directives. *Gnal*+/− mice were mated with C57BL/6J mice to produce male and female *Gnal*+/− and WT littermates. Animals (males and females *Gnal*+/− and WT aged 3 to 7-months-old) were kept at a constant room temperature and humidity on 12 h light/dark cycle and with *ad libitum* access to water and food. All the motor control experiments were performed in males and females and recordings were performed in freely moving mice.

### Open-field activity

Mice were placed in a circle arena made of plexiglass with 38 cm diameter and 15 cm height (Noldus, Netherlands) and video recorded from above. Each mouse was placed in the open-field for a period of 5 min with the experimenter out of its view. The position of centre of gravity of mice was tracked using an algorithm programmed in Python 3.5 and the OpenCV 4 library. Each frame obtained from the open-fields’ videos were analysed according to the following process: Open-field area was selected and extracted in order to be transformed into a grayscale image. Then, a binary threshold was applied on this grayscale image to differentiate the mouse from the white background. To reduce the noise induced by the recording cable or by particles potentially present in the Open-field, a bilateral filter and a Gaussian blur were sequentially applied, since those components are supposed to have a higher spatial frequency compared to the mouse. Finally, the OpenCV implementation of Canny algorithm was applied to detect the contours of the mouse, the position of the mouse was computed as mouse’s centre of mass. The distance travelled by the mouse between two consecutive frames was calculated as the variation of position of the mouse’s centre point multiplied by a scale factor, to allow the conversion from pixel unit to centimetres. The total distance travelled was obtained by summing the previously calculated distances over the course of the entire Open-field session. The speed was computed as the variation of position of centre points on two consecutive frames divided by the time between these frames (the inverse of the number of frames per seconds). This speed was then averaged by creating sliding windows of 1 second. After each session, faecal boli were removed and the floor was wiped clean with a damp cloth and dried after the passing of each mouse.

### Horizontal bar test

Motor coordination and balance were estimated with the horizontal bar test which consists of a linear horizontal bar extended between two supports (length: 90 cm, diameter: 1.5 cm, height: 40 cm from a padded surface). The mouse is placed in one of the sides of the bar and released when all four paws gripped it. The mouse must cross the bar from one side to other and latencies before falling are measured in a single trial session with a 3-min cut-off period.

### Vertical pole test

Motor coordination was estimated with the vertical pole test. The vertical pole (51 cm in length and 1.5 cm in diameter) was wrapped with white masking tape to provide a firm grip. Mice were placed heads up near the top of the pole and released when all four paws gripped the pole. The bottom section of the pole was fixated to its home-cage with the bedding present but without littermates. When placed on the pole, animals naturally tilt downward and climb down the length of the pole to reach their home cage. The time taken before going down to the home-cage with all four paws was recorded. A 20 sec habituation was performed before placing the mice at the top of the pole. The test was given in a single trial session with a 3-min cut-off period.

### Gait test

Motor coordination was also evaluated by analysing gait patterns. Mouse footprints were used to estimate foot opening angle and hindbase width, which reflects the extent of muscle loosening. The mice crossed an illuminated alley, 70 cm in length, 8 cm in width, and 16 cm in height, before entering a dark box at the end. Their hind paws were coated with nontoxic water-soluble ink and the alley floor was covered with sheets of white paper. To obtain clearly visible footprints, at least 3 trials were conducted. The footprints were then scanned and examined with the Dvrtk software (Jean-Luc Vonesch, IGBMC). The stride length was measured with hindbase width formed by the distance between the right and left hind paws.

### Grid test

The grid test is performed to measure the strength of the animal. It consists of placing the animal on a grid which tilts from a horizontal position of 0° to 180°. The animal is registered by the side and the time it drops is measured. The time limit for this experiment is 30 seconds. In those cases where the mice climbed up to the top of grid, a maximum latency of 30 seconds was applied.

### Fixed speed rotarod

Motor coordination, postural stability and fatigue were estimated with the rotarod (mouse rotarod, Ugo Basile). Facing away from the experimenter’s view, the mice placed on top of the plastic roller were tested at constant speeds (5, 10, 15, 20 and 25 r.p.m.). Latencies before falling were measured for up to 3 min in a single trial session.

### Surgery

Two surgeries were performed. During the first surgery, AAV2/1.hSyn.ChR2(H134R)-eYFP.WPRE.hGH (700nl) was injected into the deep cerebellar nuclei of the *Gnal*+/− and WT mice (dentate nucleus, DN: −6 mm AP, ±2.3 mm ML, −2.4 mm depth from dura). After 3 weeks, implantation surgery was performed. For both surgeries, the mice were anesthetized either with a mixture of ketamine/xylazine or with a mixture of isoflurane and O2 (3% for induction, 1,7% for maintenance). Injections with buprenorphine (0.05 mg/kg, s.c.) were performed to control pain, and core temperature (37°C) was maintained with a heating pad. The mice were fixed in a stereotaxic apparatus (David Kopf Instruments, USA). After a local midline lidocaine injection s.c. (2%, 1ml), a medial incision was performed exposing the skull. Small craniotomies were drilled above the recording sites and above the optic fibre location (above the virus injection site) and then the electrodes were stereotaxically lowered inside the brain. This allowed us to record in the left motor cortex (M1) (AP +2 mm and −2 mm ML from the Bregma), ventro-lateral thalamus (VAL) (−1.34 mm AP, ML=−1.00 mm and DV=-3.4 mm depth from the dura) and centro-lateral thalamus (CL) (AP at −1.58mm, ML=-0.8mm, DV=-3.00mm depth from the dura). The mice were implanted with bundles of extracellular electrodes for each recording site. The ground wire was placed on the surface of the cerebellum. Super Bond (Dental Adhesive Resin Cement, Sun Medical CO, Japan) was applied on the surface of the skull to strengthen the connection between the bone and the cement. Then, the cannulas and ground wire were fixed with dental cement (Pi-Ku-Plast HP 36, Bredent GmbH, Germany). The bundles of 8 electrodes were made in house by folding and twisting nichrome wire with 0.005 inches diameter (Kanthal RO-800) (Menardy et al., 2019). The bundles were placed inside guide cannulas (8-10 mm length and 0.16-0.18 mm inner diameter, Coopers Needle Works Limited, UK) glued (Loctite universal glue) to an electrode interface board (EIB-16; Neuralynx, Bozeman, MT, USA) with 1 wire for each channel and 4 channels for each brain region (M1, CL, VAL), extending 0.5 mm below the tube tip. Wires were then fixed to the EIB with gold pins (Neuralynx, Bozeman, MT, USA) and then the EIB was secured in place by dental cement. A gold solution (cyanure-free gold solution, Sifco, France) was used for gold plating and the impedance of each electrode was set to 200–500 kΩ.

### Manipulations of cerebellar output

Because the cerebellar nuclei sends projections in the contralateral thalamus that then connects with the M1 (Teune et al., 2004) optogenetic stimulations were performed in the contralateral cerebellar DN (left M1 and right DN). Light-induced excitation of the cerebellar-projection neurons was elicited by using a LED driver (Mightex Systems^®^) through optical fibres radiating blue light (470 nm) unilaterally implanted into the deep cerebellar nuclei (light intensity of ~1.5mW/mm2). Optogenetic stimulations of the DN (1000 mA, 100ms, 0.25Hz or theta-burst stimulation 1000 mA, 20ms, 8,33Hz, applied for 2 × 40s with a 2 min pause in between) were performed before and after triggering the dystonic attacks by an oxotremorine methiodide (oxotremorine M) intraperitoneal injection.

### Electrophysiological recordings

The recordings begun after at least 3 days of recovery and were performed on awake freely moving mice using a 16-channel acquisition system with a sampling rate of 25 kHz (Tucker Davis Technology System 3, Tucker-Davis Technologies, Alachua, FL, USA). We performed 60 minutes baseline recording in an open-field, followed by a 60 min recording after a saline injection. *Gnal*+/− and WT mice were then injected intraperitoneally with oxotremorine methiodide (0.1 mg/kg, Sigma-Aldrich), dissolved in saline (NaCl 0.9g/L) and recorded another 60 min with the same protocol as for saline. Optogenetic stimuli were applied on the DN at low frequency stimuli of 100 ms, 1000 mA and 0.25 Hz. On the next day, theta burst stimulations was implemented. The mice were recorded 60 min after the saline injection and then 60 min after the oxotremorine M injection using the same protocol as in baseline day except that after 30 minutes 2 theta-burst sessions of 40 sec each with 2 minutes pause in between were applied (a total of 600 pulses for each condition).

### Histological verification of the site of optical fibre in DN and verification of the position of the electrodes

The animals were sacrificed with a single dose of pentobarbital (100mg/kg, i.p.). Electrolytic lesions were performed to check the position of the electrodes, mice were perfused with paraformaldehyde and the brains were removed and kept in paraformaldehyde (4%). After slicing (using a vibratome at 90 μm thickness), all sites of the recordings were verified by superposing the atlas (Allen Brain Atlas) on slices, with the closest anatomical landmarks from our lesions used as reference points. *(Figure 1.D)*

### Behavioural Analysis

Video recordings monitored the motor behaviour of *Gnal*+/− and WT mice in the open-field. Dystonia severity was estimated using a previously published abnormal movement scoring scale (Jinnah et al., 2000; Calderon et al., 2011; Pelosi et al., 2017) for every 10 min-long block of the recording, after the oxotremorine M injection was performed. The assessment was blinded for mouse genotype and was done by 2 members of the team. The scale uses the following scores: 0= normal motor behavior; 1=no impairment, but slightly slowed movements; 2. Mild impairment: occasional abnormal postures and movements; ambulation with slow walk; 3. Moderate impairment: frequent abnormal postures and movements with limited ambulation; 4. Severe impairment: sustained abnormal postures without any ambulation or upright position. In addition, total time of active wakefulness from the total time of recording (active wake percentages, AW%) was assessed for both states, pre and post oxotremorine and pre and post theta-burst stimulation. Active wake was considered as the state when the mouse is exploring the open-field by walking in any direction and is expressed as a percentage of the total time of the recording (Georgescu et al., 2018). We evaluated the impact of theta-burst DN stimulation on the onset of dystonic-like symptoms in saline- and oxotremorine-treated *Gnal*+/− mice.

### Electrophysiological Analysis

Spike sorting was completed using homemade Matlab scripts (Mathworks, Natick, MA, USA) that are based on *k-means clustering on PCA of the spike waveforms* (Paz et al., 2006). In order to evaluate the activity of the same cells in similar conditions during experiments, we investigated the change of the firing rate probability in the thalamus and M1 motor cortex during cerebellar DN 100ms stimulations after saline or oxotremorine M administration performed and analyses in one continuous session. The average increase in firing rate during the stimulation was determined by computing the peri-stimulus time histogram (bin: 10ms) of the spikes around the stimulation; the spike count in the histogram was divided by the duration of the stimulation and the number of stimulations administered to yield a firing rate. The acceleration of discharge due to the stimulation was taken as the average spike count during the stimulation subtracted by the baseline (taken as the 300ms that preceded the stimulation onset). The response to stimulation was only analysed in cells where at least one bin during the stimulation was larger than 4 times the standard deviation of the baseline values

### Statistics

Figures represent the averages ± standard error of the mean (SEM). Student T test, Mann-Whitney test, repeated-measure ANOVA and multiple comparisons test was used as appropriate, after testing the normality of the data by D’Agostino and Pearson omnibus normality test.

